# Validating static food web model diet matrices using stable isotope simulation

**DOI:** 10.64898/2026.10.30.762321

**Authors:** Megan J. Siemann, Kieran M. Tierney, William F. Patterson, David Chagaris

**Affiliations:** School of Natural Resources and Environment, University of Florida, Gainesville, FL, USA; Nature Coast Biological Station, University of Florida, Gainesville, FL, USA; SUERC, University of Glasgow, Scottish Enterprise Technology Park, East Kilbride, UK; Forestry, Fisheries, and Geomatic Sciences, University of Florida, Gainesville, FL USA

**Keywords:** Ecopath, stable isotope, carbon, nitrogen, sulfur

## Abstract

End-to-end ecosystem models, such as Ecopath with Ecosim (EwE), are valuable tools for resource management. Within these frameworks, trophic interactions, typically derived from stomach contents analyses, govern system-wide energy flow via consumption. However, stomach contents data may be unrepresentative of long-term average diets and assimilated consumption due to factors including differential digestion rates and identifiability of prey, introducing model uncertainty. Because trophic interactions define model structure, variation in modeled diets can alter predictions and management conclusions. Consequently, evaluating and reducing uncertainty in diet matrix compositions is critical for operational acceptance of ecosystem models. Stable isotope analysis provides complementary, time-integrated insight into average diets, but its application for EwE validation has largely been limited to evaluating trophic levels and isotopic niche space. Here, we present a quantitative framework for simulating consumer stable isotope signatures across food webs from parent ecosystem models using a matrix-based mixing model. Using a published Northern Gulf of Mexico EwE model as a case study, we demonstrate Monte Carlo simulation to propagate uncertainty from basal source signatures and trophic discrimination factors. Comparison of predicted signatures with empirical δ¹³C and δ³⁴S measurements enables validation of primary carbon sources and benthic–pelagic pathway contributions, revealing agreement in isotope space alongside mismatch in energy pathways. Diagnostic residuals identify discrepancies in diet composition. Model scoring provides a reproducible framework for ranking alternative diet configurations. Redundancy analysis further identifies within-group species coverage differences between stomach contents and isotope datasets as a primary driver of mismatch, highlighting an important consideration for sampling design.

## 1. Introduction

Ecosystem modeling has become an essential component of ecosystem-based management (EBM), providing critical tools for understanding energy fluxes and mapping macro-trophic pathways within complex biological networks (Heymans et al. 2016; Rodriguez-Perez et al. 2023). Characterizing these energy dynamics allows resource managers to comprehend ecosystem stability, predict structural shifts, and identify potential systemic vulnerabilities (Rodriguez-Perez et al. 2023). Within the food web, ambient energy from the environment is channeled through production and consumption: primary producers capture solar energy via photosynthesis, and this captured energy is subsequently transmitted through higher trophic levels via predator-prey interactions. End-to-end ecosystem models incorporate this trophic energy flow by explicitly defining foraging dynamics and predator-prey relationships within their mathematical frameworks (Christensen and Walters 2004a; Fulton et al. 2011). Although model development can require substantial data and time, once established, end-to-end ecosystem models are versatile tools for evaluating the impacts of management decisions, natural disturbances, and environmental change (Fulton et al. 2011).

The increasing use of end-to-end ecosystem models has driven advances in model capability, alongside a need for improved validation and quantification of uncertainty. Frameworks such as Ecopath with Ecosim (EwE) (Christensen and Walters 2004b) and Atlantis (Fulton et al. 2011) are widely applied in marine systems. EwE, originally developed to help fisheries managers understand ecosystem structure (Polovina 1984), has evolved into a dynamic framework capable of projecting biomass, consumption, and energy flows across time and space under alternative environmental and management scenarios (Pauly et al. 2000; Colleter et al. 2015; Villasante et al. 2016; de Mutsert et al. 2024). However, differences in model parameterization, particularly of trophic dynamics, can strongly influence model predictions, and uncertainty remains a barrier to broader adoption (Karp et al. 2023). Confidence in the diet matrix is especially critical, as trophic relationships form a core structural component of these models (Ainsworth et al. 2010).

Consequently, improving diet matrix parameterization and validation, through both methodological advances and expanded data sources, remains a key priority (Anstead et al. 2021; Karp et al. 2023).

### 1.1 Diet matrix challenges

Trophic interactions in food web models are defined by a diet matrix, in which diet compositions are expressed as proportions of prey items for each consumer and must sum to one to maintain mass balance (Christensen and Walters 2004a). Historically, stomach contents analysis (SCA) provided the basis for estimating these proportions. However, SCA is subject to several limitations, including high occurrence of empty stomachs, differential digestion rates, and the presence of cryptic or unidentifiable prey items (Amundsen and Sánchez-Hernández 2019). Some of these challenges can be mitigated through increased sampling effort, but inherent biases remain. For example, soft-bodied prey degrade more rapidly and are less likely to be identified than prey with hard parts, leading to potential underrepresentation of certain prey types (Dahl et al. 2017; Nielsen et al. 2018). Additionally, species with diverse diets may require extensive sampling to adequately characterize their feeding habits (Vinson and Budy 2011).

These limitations contribute to substantial uncertainty in diet matrix parameterization (Trijoulet et al. 2019). Data constraints further complicate this process, as diet information is often compiled from multiple studies or repositories such as FishBase (Froese and Pauly 2026) and the Gulf of Mexico Species Interactions Database (GoMexSI; Simons et al. 2014). Region-and time-specific data may be unavailable even in well-studied systems, necessitating the use of “best available” data from analogous systems or species (Dowling et al. 2023). Although best-practice guidelines exist (Heymans et al. 2016; Dowling et al. 2023), parameterization is ultimately constrained by data availability. These challenges have motivated the development of alternative approaches for diet estimation and for quantifying associated uncertainty.

Error in diet composition estimates can lead to misrepresentation of energy flow and, consequently, biased model outputs (Ainsworth et al. 2010; Masi et al. 2014). To address these issues, researchers have increasingly incorporated additional trophic data sources. Molecular techniques, including DNA barcoding, improve identification of highly digested or cryptic prey items (Dahl et al. 2017; Nielsen et al. 2018), while stable isotope analysis provides time-integrated measures of diet (Hobson and Clark 1992; Vander Zanden et al. 2015). Each method contributes complementary information, with DNA-based approaches offering high taxonomic resolution and stable isotopes capturing longer-term dietary integration (Nielsen et al. 2018).

Stable isotope ratios in consumer tissues reflect those of their food sources, as assimilated material is incorporated into tissues over time (Peterson and Fry 1987; Hobson and Clark 1992). Lighter isotopes are preferentially metabolized, resulting in enrichment of heavier isotopes in consumer tissues. The predictable enrichment of approximately 3.5‰ per trophic level, known as the trophic discrimination factor (TDF), makes δ^15^N a useful tracer of trophic position (Post 2002). In contrast, δ^13^C exhibits lower fractionation (∼1.0‰ per trophic level) but varies among primary producers due to differences in photosynthetic pathways, allowing inference of basal carbon sources (Peterson and Fry 1987; Post et al. 2007). δ^34^S can further distinguish between benthic and pelagic pathways in some aquatic or marine ecosystems due to differences generated by sulfate reduction in sediments, although its application has been more limited (Raoult et al. 2024). Together, these tracers provide valuable insight into trophic niches and energetic pathways.

### 1.2 Stable isotopes and trophic validation

Diet estimation methods have evolved from deterministic averaging toward statistical approaches that explicitly incorporate uncertainty (Masi et al. 2014) Monte Carlo resampling techniques have been implemented within EwE through tools such as Ecoranger and Ecosampler (Morissette 2005; Steenbeek et al. 2018), enabling exploration of diet matrix uncertainty. Stable isotopes have also been incorporated into diet estimation, as their time-integrated nature can better represent average diet and mitigate some biases associated with SCA (Nielsen et al. 2018; Trijoulet et al. 2019; Hernvann et al. 2022). Stable isotope mixing models estimate the proportional contributions of prey sources based on observed isotope signatures, typically δ^13^C and δ^15^N. These models can incorporate prior information, including SCA-derived prey likelihoods, but are generally applied at the level of individual consumers rather than across entire food webs (Polito et al. 2011; Giménez et al. 2017). Bayesian approaches have further extended this framework to integrate multiple data sources (Hernvann et al. 2022; Lobry et al. 2022). Despite these advances, uncertainty in diet composition remains.

Stable isotopes have also been used to evaluate food web models. Previous validation efforts have compared model-derived metrics with isotope-based estimates, rather than directly comparing predicted and observed isotope signatures. For example, EwE-derived trophic levels have been compared with δ^15^N-based estimates to assess trophic structure (Kline Jr and Pauly 1998; Navarro et al. 2011; Deehr et al. 2014; Pedersen 2022). Similarly, δ^13^C and δ^15^N have been used to estimate isotopic niche width, which has been compared with the EwE omnivory index (Navarro et al. 2011; Ramsvatn 2013).

Some authors have explored benthic versus pelagic energy contributions using isotope-derived proxy metrics; however, these approaches have not directly predicted stable isotope signatures from model consumption flows and have been limited to δ^13^C and δ^15^N (Wang et al. 2020). The use of δ^34^S for model evaluation remains largely unexplored (Raoult et al. 2024).

Importantly, existing validation approaches rely on derived metrics rather than testing whether modeled consumption flows are consistent with observed isotope data. To our knowledge, there is currently no published method to evaluate whether diet matrix parameterization produces isotope signatures consistent with empirical observations or accurately represents basal-source and benthic–pelagic energy pathways. A forward isotope simulation approach therefore offers a means to directly link the diet matrix of a static Ecopath base model with observed isotope data, providing an additional diagnostic layer for ecosystem model validation.

### 1.3 Objectives

The overarching goal of this work is to improve our ability to validate trophic-dynamic ecosystem models through direct comparison with empirical stable isotope data. In this study, we introduce a forward stable isotope simulation framework that predicts consumer isotope ratios from diet proportions specified in Ecopath diet matrices and compares these predictions with empirical data. We demonstrate this framework using the Northern Gulf of Mexico EwE model (Chagaris et al. 2020) and site-specific stable isotope data (Dahl and Patterson 2014; Tarnecki and Patterson 2015; Patterson III et al. 2020) to show how it can be used to diagnose mismatches and evaluate alternative diet matrix configurations.

## 2. Materials and Methods

### 2.1 Framework overview

We developed a forward simulation framework that predicts consumer stable isotope signatures from diet proportions specified in a parent food web model to address uncertainty in trophic interactions and evaluate diet matrix parameterization (Figure 1). In contrast to traditional isotope mixing models, which are inverse approaches that estimate average diet from stable isotope signatures integrated over time in consumer tissues (Smith et al. 2013; Stock et al. 2018), this framework uses proportional trophic flows as fixed inputs and computes the isotope values implied by those flows. Uncertainty in basal-source isotope signatures and trophic discrimination factors (TDFs) is propagated using Monte Carlo simulation, generating predictive probabilistic distributions for consumer isotope values. Model performance is evaluated by comparing predicted values with empirical stable isotope data, enabling diagnostic assessment of trophic structure and diet matrix specification.

**Figure 1.**
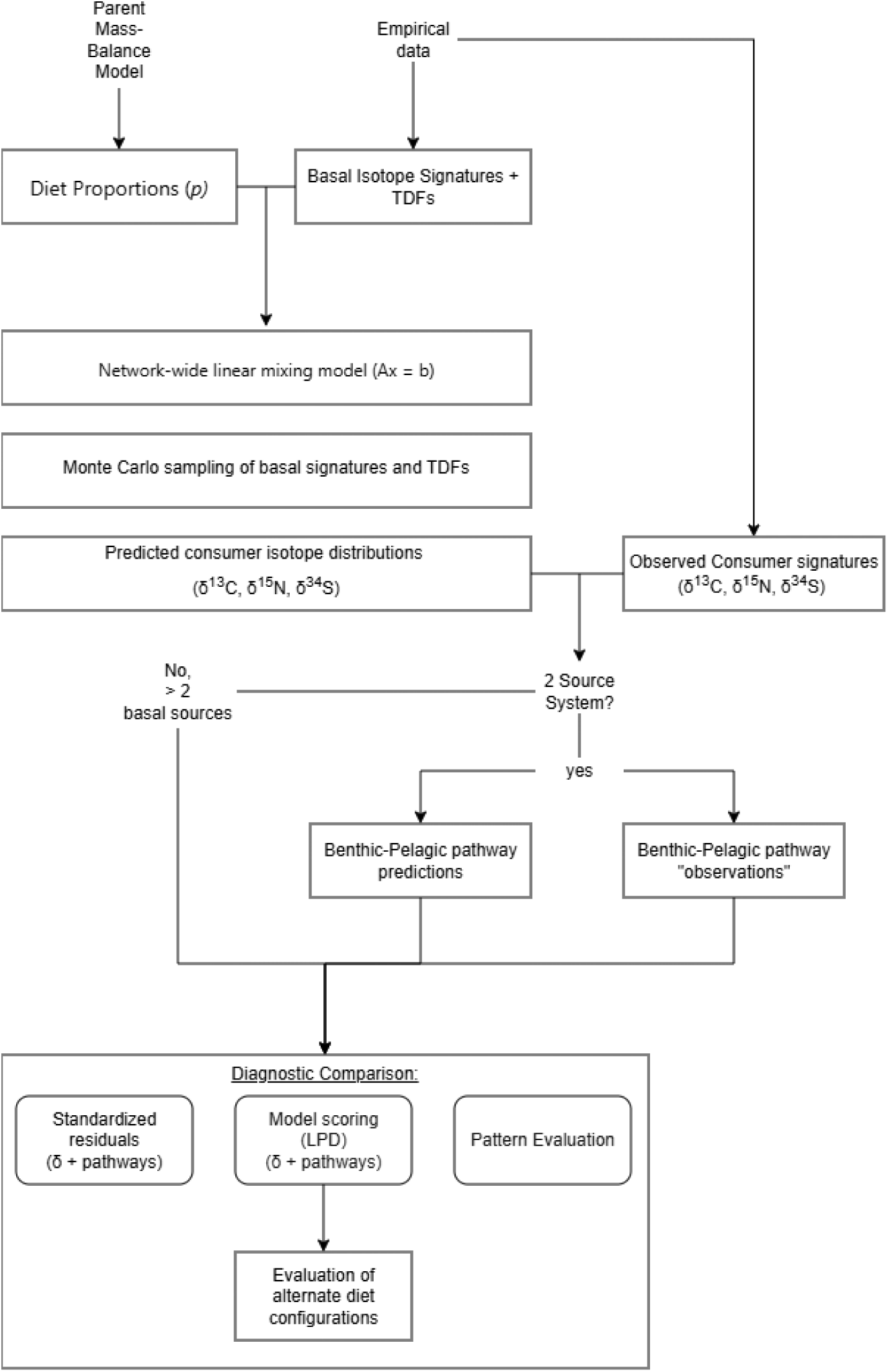
Workflow of the forward fish muscle stable isotope simulation framework, showing propagation of diet matrices, basal signatures, and trophic discrimination factors through a network-wide mixing model to generate predicted consumer isotope signatures and diagnostic comparisons.

#### 2.1.1 Network-wide isotope prediction

The following inputs are required for the forward mixing model:

1. Stable isotope values for all basal sources (*s*)

2. Diet proportions (*p*) from the parent mass-balance food web model

3. TDFs for each isotope (*k*) for the system or by consumer (*j*).

Additional empirical consumer isotope data are required for model evaluation.

Primary producers and detrital groups are treated as basal sources in the model. Detrital material undergoes further biogeochemical transformation (e.g., remineralization and nutrient recycling), which can alter isotopic signatures from freshly aggregated detritus. To avoid propagation of these discrepancies through the system, detrital groups are treated as independent basal sources.

For a food web with *S* basal groups and *J* consumer groups, the total number of groups is *G* = *S* + *J*. All groups may contribute as a dietary source and therefore act as prey (*i*).Consumer isotope values are modeled as weighted sums of prey isotope values plus a trophic discrimination factor, consistent with standard mixing models (Stock et al. 2018; Equation 1): For clarity, Equation 1 is shown for a single isotope and consumer. In subsequent equations, isotope-specific notation is introduced and the formulation extended to the full food web.

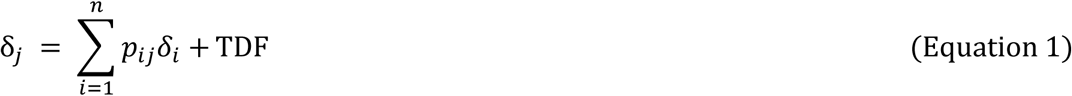

This formulation is rearranged into a linear system (Equation 2–3). The system is solved independently for each isotope *k* as *Ax_k_* = *b_k_*, where *A* contains the food-web structure and *x_k_*and *b_k_* contain isotope-specific values and parameters.

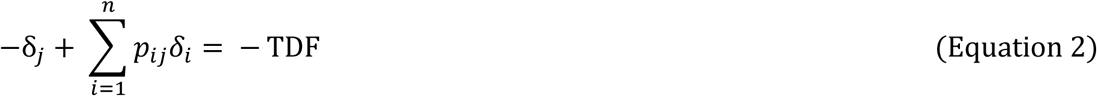

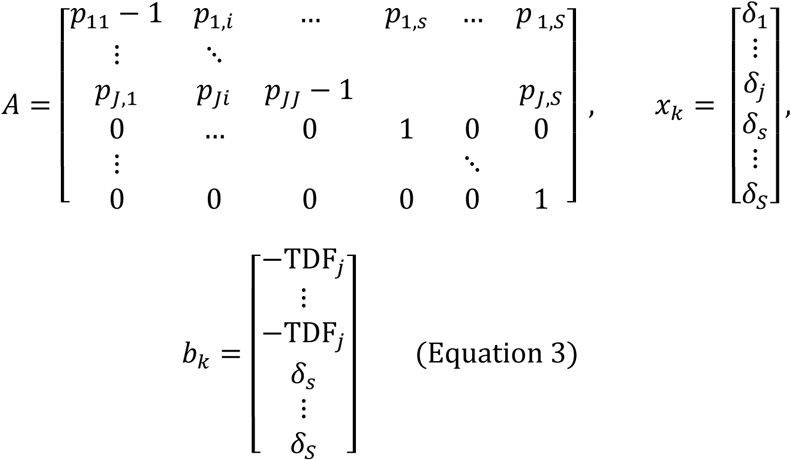

The linear system in Equation 3 compactly represents the full network of isotope mixing relationships across all groups. The matrix *A* contains the diet structure of the food web, *x_k_* is the vector of isotope signatures for all groups, and *b* is the vector of constants defined by trophic discrimination factors (TDFs) for consumers and fixed isotope values for basal sources.

Each consumer row in *A* corresponds to the mixing equation for a single consumer. Off-diagonal elements in that row, *A_ji_*, are equal to the proportional contribution of prey group *i* to consumer *j* (i.e., *P_ji_*). The diagonal element, *A_jj_*, is equal to −1 + *p_jj_*, which accounts for any cannibalistic contribution. For consumers without cannibalism, this reduces to −1. As a result, each consumer row sums to zero, reflecting conservation of mass in the mixing relationship. The corresponding entry in the vector *b_k_* for each consumer row is the negative trophic discrimination factor, −TDF*_j,k_*, for isotope *k*. This term represents the enrichment between the consumer and its diet. Each basal group row imposes a fixed boundary condition. For these rows, the diagonal element of *A* is set to 1 and all other entries are 0, meaning that the isotope value of the basal group is fixed and does not depend on other groups. The corresponding entry in *b* is simply the observed isotope value of that basal source.

Solving the system *Ax = b* therefore yields a set of isotope values *x =* (*δ_1_*,…, *δ_n_*) that are internally consistent with the specified diet matrix, basal isotope values, and trophic discrimination factors across the entire food web.

#### 2.1.2 Monte Carlo simulation

The linear model produces a single deterministic solution for consumer isotope values. To account for uncertainty, Monte Carlo (MC) simulation is used to repeatedly sample input parameters and generate predictive distributions of isotope values. In each iteration (*m*), basal-source isotope signatures and TDFs *δ_s,k_*, and *TDF_j,k_*, are sampled from predefined distributions, and the forward mixing model is solved. This produces distributions of predicted consumer isotope signatures, *δ_j,k_*, across simulations. Diet proportions are not varied within the MC draws due to the high dimensionality of the diet matrix. Instead, variability in diet composition is assessed separately through diagnostic residuals and model comparison procedures.

#### 2.1.3 Benthic–pelagic pathway contributions

As a derived diagnostic, benthic-pelagic pathway contributions were estimated from the predicted isotope values generated by the forward model. For systems with two distinct basal end members, benthic and pelagic pathway contributions can be estimated. Similar approaches have been used previously to estimate pathway utilization (Kopp et al. 2015; von Biela et al. 2016; Wang et al. 2022). In this framework, pathway contributions are derived from predicted isotope values using trophic-adjusted endpoints (Equations 4–6).

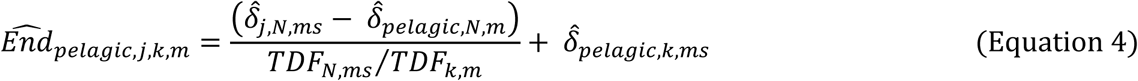

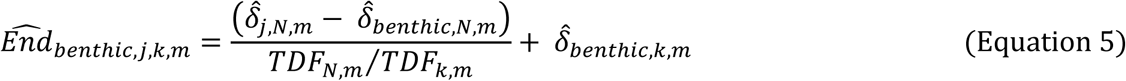

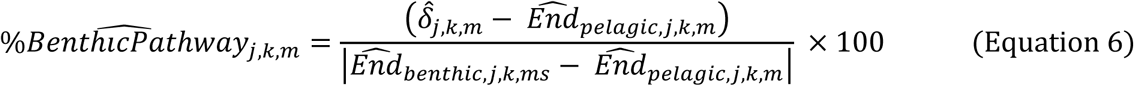

Nitrogen (*N*) is used to account for trophic position, and the ratio of TDFs is used to scale adjustments. Observed pathway contributions are calculated separately using mean TDFs and basal-source signatures (Equations 7–9) to avoid instability from extreme MC draws.

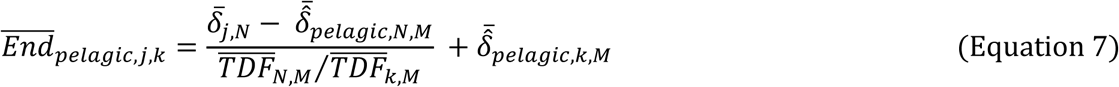

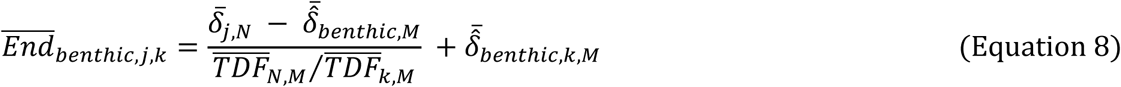

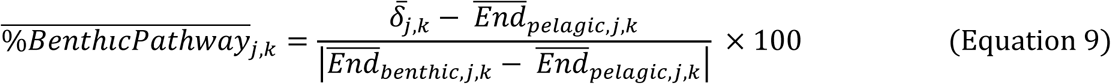

#### 2.1.4 Diagnostic comparison and residuals

Parent model diet representation is evaluated by comparing predicted and observed isotope values (δ¹⁵N, δ¹³C, δ³⁴S). Standardized residuals are calculated as the difference between observed and predicted values, scaled by combined observational and predictive uncertainty. Residuals are computed for both directly predicted isotope values and for derived benthic–pelagic pathway contributions. These residuals quantify the magnitude and direction of mismatch and are used diagnostically to identify patterns of disagreement between isotope-implied diet and the parent model diet representation across consumers and tracers.

#### 2.1.5 Model Scoring

Model scoring procedures allow for objective comparison of how alternative trophic configurations agree with observed stable isotope data. Variability in modeled diet composition was evaluated using alternative diet matrices derived either from alternative estimation methods or from stochastic variation in prey proportions. Alternate diet matrices are evaluated using log predictive density (LPD) (Gelman et al. 2014) which quantifies the likelihood of observed isotope data under model-predicted distributions. For each consumer group and isotope, predicted values from the Monte Carlo distribution were evaluated against the fitted observed-data distribution, and pointwise log densities were summed across groups. Higher LPD values indicate better agreement between modeled and observed isotope signatures. This provides a reproducible framework for comparing alternative trophic configurations.

### 2.2 Northern Gulf of Mexico case study

#### 2.2.1 Northern Gulf of Mexico EwE model

An existing, calibrated Ecopath with Ecosim (EwE) model of the Northern Gulf of Mexico (NGoM-EwE) (Chagaris et al. 2020) was used to demonstrate the stable isotope simulation and diet matrix validation framework. The NGoM-EwE model was developed to examine the relative effects of multiple stressors in the system, including the 2010 Deepwater Horizon oil spill (DWH) and the expansion of invasive lionfish (*Pterois volitans*) (Dahl and Patterson 2014; Tarnecki and Patterson 2015; Dahl et al. 2017; Lewis et al. 2020; Patterson III et al. 2020). The model represents a reef-associated food web with strong benthic–pelagic coupling and includes two detrital groups, four primary producer groups, and 58 consumer groups. Commercially and ecologically important species are resolved at the species level, often with age stanzas, while remaining taxa are aggregated into functional groups based on trophic role and ecological similarity. Phytoplankton and benthic primary producers account for basal production, with detritus providing an additional supporting pool. Although seagrass is included, benthic production is dominated by macroalgae and microphytobenthos, both in the model (Chagaris et al. 2020) and in the shelf ecosystem (Dahl and Patterson 2014; Tarnecki and Patterson 2015). This results in distinct benthic and pelagic signatures for both δ¹³C and δ³⁴S (Dahl and Patterson 2014; Tarnecki and Patterson 2015). The presence of well-separated basal isotope sources, combined with paired stomach contents and isotope data, makes the NGoM-EwE model well suited for evaluating energy pathway representation.

The diet matrix for the published model was parameterized using locally collected stomach contents data (Dahl and Patterson 2014; Tarnecki and Patterson 2015) and supplemented with additional datasets, including Southeast Area Monitoring and Assessment Program (SEAMAP), Florida Fish and Wildlife Conservation Commission (FWC), GoMexSI, and FishBase data.

Paired stomach contents and stable isotope data were available for many samples, providing site-specific empirical constraints. Diet proportions for invertebrate groups were not specifically estimated for the NGoM-EwE model but were instead borrowed from the West Florida Shelf EwE model (Okey et al. 2004; Chagaris et al. 2015, 2017). The balanced diet matrix from the published model was used as the baseline (“published”) configuration for comparison.

Alternative diet matrices were developed using the aggregated SCA data to evaluate sensitivity to diet parameterization and demonstrate model scoring. These included pre-balance diet proportions estimated using weighted averages and maximum likelihood estimates based on a Dirichlet–multinomial formulation (Masi et al. 2014). These matrices enabled evaluation of major differences arising from alternative diet estimation methods, as well as the effects of adjustments introduced during Ecopath model balancing. EwE allows for inclusion of an imported prey category to account for consumption outside of the model scope or groups defined within. For all diet matrices, any imported prey proportions were excluded and remaining prey items re-summed to 1 for mass balance.

To further explore variability in diet composition within empirically supported bounds, 10,000 stochastic realizations were generated by sampling from Dirichlet distributions constrained by 95% confidence intervals of prey proportions derived from the Dirichlet–multinomial formulation. Within these randomized diet matrices, prey items could be excluded when supported by the bounds, allowing for diet simplification and diets of invertebrate groups lacking SCA data were varied within pooled confidence intervals (Supplementary Figure 1).

#### 2.2.2 Observed consumer stable isotope data

Analyses were based on stable isotope ratios measured from muscle tissue, which integrates assimilated dietary signals over periods of months and therefore provides a time-averaged representation of trophic interactions appropriate for comparison with Ecopath diet matrices (Hobson and Clark 1992; Vander Zanden et al. 2015). Observed consumer isotope values (δ¹⁵N, δ¹³C, and δ³⁴S) were compiled from unpublished data (W. Patterson) and previously published datasets (Dahl and Patterson 2014; Tarnecki and Patterson 2015; Patterson III et al. 2020).

Muscle tissue samples were collected, dried, and processed following established protocols prior to mass spectrometry analysis. δ¹³C values were lipid-corrected for samples with C:N ratios greater than 3.5 (Post et al. 2007). Group-level isotope data were summarized using location–scale t-distributions to reflect observed variability and to provide likelihood functions for model evaluation (Table 1).

**Table 1.** Observed fish muscle stable isotope data and fitted location-scale t-distribution parameters. Scientific names are provided for single species groups.

| Group<br># | Group name,<br>*Scientific Name* | Trophic<br>Level | δ<sup>13</sup>C |  |  |  |  | δ<sup>15</sup>N |  |  |  |  | δ<sup>34</sup>S |  |  |  |  |
|---|---|---|---|---|---|---|---|---|---|---|---|---|---|---|---|---|---|
|  |  |  | Observed |  |  | t distribution |  | Observed |  |  | t distribution |  | Observed |  |  | t distribution |  |
|  |  |  | *n* | μ | *SD* | *v* | σ | *n* | μ | *SD* | *v* | σ | *n* | μ | *SD* | *v* | σ |
| 1 | pelagic sharks | 4.31 | 1 | -16.98 | . | . | . | 1 | 15.16 | . | . | . | 1 | 16.22 | . | . | . |
| 3 | rays and small sharks* | 3.78 | 3 | -17.06 | 0.30 | 7 | 0.64 | 3 | 13.18 | 0.19 | 7 | 0.96 | 3 | 16.38 | 1.16 | 7 | 1.16 |
| 4 | large pelagic piscivores | 4.26 | 38 | -17.45 | 0.65 | 7 | 0.55 | 38 | 13.47 | 1.11 | 7 | 0.94 | 35 | 17.98 | 1.06 | 7 | 0.90 |
| 5 | greater amberjack<br>*Seriola dumerili* | 4.38 | 41 | -16.70 | 0.39 | 7 | 0.33 | 41 | 14.11 | 1.00 | 7 | 0.84 | 38 | 17.55 | 0.81 | 7 | 0.68 |
| 6 | small pelagics | 3.05 | 36 | -18.29 | 0.53 | 7 | 0.44 | 36 | 12.31 | 1.50 | 7 | 1.27 | 35 | 18.56 | 0.96 | 7 | 0.81 |
| 7 | reef carnivores | 4.07 | 70 | -17.36 | 0.39 | 7 | 0.33 | 70 | 13.76 | 0.82 | 7 | 0.69 | 65 | 18.95 | 0.99 | 7 | 0.84 |
| 9 | gag 1-2\*<br>*Mycteroperca microlepis* | 4.27 | 2 | -17.05 | 0.20 | 7 | 0.64 | 2 | 14.10 | 0.17 | 7 | 0.96 | 2 | 17.49 | 0.85 | 7 | 1.16 |
| 10 | gag 3+<br>*Mycteroperca microlepis* | 4.29 | 9 | -16.78 | 0.50 | 7 | 0.42 | 9 | 14.37 | 0.95 | 7 | 0.81 | 7 | 16.23 | 1.20 | 7 | 1.01 |
| 12 | red grouper 1-2<br>*Epinephelus morio* | 3.97 | 1 | -16.70 | . | . | . | 1 | 13.12 | . | . | . | 1 | 18.05 | . | . | . |
| 13 | red grouper 3+<br>*Epinephelus morio* | 4.41 | 23 | -16.65 | 0.23 | 7 | 0.20 | 23 | 13.59 | 0.83 | 7 | 0.70 | 20 | 17.97 | 0.64 | 7 | 0.54 |
| 14 | red snapper juvenile<br>*Lutjanus campechanus* | 4.03 | 179 | -17.29 | 0.47 | 7 | 0.39 | 179 | 13.72 | 0.87 | 7 | 0.73 | 158 | 18.48 | 1.13 | 7 | 0.95 |
| 15 | red snapper adult<br>*Lutjanus campechanus* | 4.09 | 530 | -17.30 | 0.46 | 7 | 0.39 | 529 | 13.68 | 0.75 | 7 | 0.63 | 439 | 18.35 | 1.33 | 7 | 1.13 |
| 16 | gray snapper<br>*Lutjanus griseus* | 3.89 | 9 | -16.54 | 0.37 | 7 | 0.32 | 9 | 14.06 | 0.47 | 7 | 0.39 | 9 | 17.66 | 1.06 | 7 | 0.89 |
| 17 | lane snapper<br>*Lutjanus synagris* | 3.66 | 20 | -16.72 | 0.70 | 7 | 0.59 | 20 | 13.70 | 0.50 | 7 | 0.43 | 19 | 18.09 | 0.95 | 7 | 0.80 |
| 18 | gray triggerfish<br>*Balistes capriscus* | 3.81 | 287 | -17.99 | 0.55 | 7 | 0.46 | 287 | 12.39 | 0.85 | 7 | 0.72 | 218 | 18.64 | 1.61 | 7 | 1.36 |
| 19 | reef crustacean eaters | 3.74 | 107 | -17.54 | 0.67 | 7 | 0.57 | 107 | 12.15 | 1.23 | 7 | 1.04 | 103 | 17.26 | 2.06 | 7 | 1.74 |
| 20 | reef invertivores | 3.31 | 17 | -18.26 | 0.64 | 7 | 0.54 | 17 | 11.70 | 1.29 | 7 | 1.09 | 17 | 18.98 | 0.47 | 7 | 0.39 |
| 21 | medium reef omnivores | 3.36 | 7 | -17.91 | 0.41 | 7 | 0.35 | 7 | 12.55 | 1.21 | 7 | 1.03 | 7 | 18.46 | 1.16 | 7 | 0.98 |
| 22 | small reef omnivores | 3.14 | 7 | -19.18 | 1.37 | 7 | 1.16 | 7 | 14.58 | 0.44 | 7 | 0.37 | 7 | 16.89 | 1.45 | 7 | 1.23 |
| 23 | reef planktivores | 3.34 | 19 | -17.82 | 0.46 | 7 | 0.39 | 19 | 12.32 | 0.66 | 7 | 0.55 | 17 | 18.89 | 0.62 | 7 | 0.52 |
| 24 | porgies | 3.68 | 126 | -17.24 | 0.45 | 7 | 0.38 | 126 | 13.07 | 0.57 | 7 | 0.48 | 86 | 17.81 | 1.04 | 7 | 0.88 |
| 25 | grunts | 3.51 | 152 | -17.27 | 0.55 | 7 | 0.46 | 152 | 12.76 | 0.61 | 7 | 0.51 | 107 | 16.97 | 2.03 | 7 | 1.72 |
| 26 | lionfish juv<br>*Pterois volitans* | 3.89 | 62 | -17.86 | 0.46 | 7 | 0.39 | 54 | 12.43 | 0.75 | 7 | 0.63 | 54 | 19.19 | 0.52 | 7 | 0.44 |
| 27 | lionfish adult<br>*Pterois volitans* | 4.35 | 240 | -17.71 | 0.34 | 7 | 0.29 | 208 | 12.70 | 0.62 | 7 | 0.52 | 208 | 19.35 | 0.61 | 7 | 0.51 |
| 29 | small wrasses | 3.61 | 14 | -18.60 | 0.70 | 7 | 0.59 | 14 | 11.65 | 1.17 | 7 | 0.99 | 12 | 19.57 | 0.82 | 7 | 0.69 |
| 30 | vermilion snapper<br>*Rhomboplites aurorubens* | 3.65 | 248 | -18.27 | 0.44 | 7 | 0.37 | 248 | 12.78 | 0.67 | 7 | 0.57 | 186 | 19.17 | 0.74 | 7 | 0.62 |
| 31 | chromis | 3.20 | 8 | -18.69 | 0.36 | 7 | 0.30 | 8 | 11.32 | 0.77 | 7 | 0.65 | 8 | 20.39 | 0.18 | 7 | 0.15 |
| 32 | damselfish | 2.67 | 8 | -18.30 | 0.65 | 7 | 0.55 | 8 | 11.76 | 0.29 | 7 | 0.24 | 8 | 18.99 | 0.98 | 7 | 0.83 |
| 33 | seabasses | 3.52 | 99 | -17.47 | 0.51 | 7 | 0.43 | 99 | 12.46 | 0.92 | 7 | 0.78 | 81 | 18.17 | 0.94 | 7 | 0.79 |
| 34 | sand perch | 3.87 | 51 | -17.48 | 0.55 | 7 | 0.47 | 51 | 11.92 | 1.18 | 7 | 1.00 | 39 | 16.91 | 0.88 | 7 | 0.74 |
| 35 | lizardfish | 4.05 | 51 | -17.89 | 0.42 | 7 | 0.36 | 51 | 12.47 | 1.40 | 7 | 1.19 | 51 | 17.70 | 1.43 | 7 | 1.21 |
| 37 | demersal carnivores | 3.76 | 9 | -19.04 | 1.47 | 7 | 1.24 | 9 | 13.39 | 1.95 | 7 | 1.64 | 9 | 14.86 | 3.32 | 7 | 2.81 |
| 38 | demersal crustacean eaters | 3.56 | 63 | -18.09 | 1.59 | 7 | 1.34 | 63 | 12.55 | 1.22 | 7 | 1.03 | 63 | 16.25 | 3.55 | 7 | 3.00 |
| 39 | demersal omnivores | 3.15 | 105 | -17.91 | 1.60 | 7 | 1.35 | 105 | 12.82 | 1.42 | 7 | 1.20 | 105 | 15.14 | 2.73 | 7 | 2.31 |

\* marked groups assigned pooled sigma value

Samples were restricted to the spatial extent of the NGoM-EwE model (shoreline to 29°N and −88.5° to −85.5°) to account for spatial variability in isotopic baselines (Radabaugh et al. 2013). Although the Ecopath base model represents conditions in 2009, isotope data collected between summer 2009 and fall 2015 were retained to maximize coverage of consumer groups. The final dataset included 34 of the 58 consumer groups in the model.

#### 2.2.3 Parameterization and simulation of isotope predictions

Stable isotope predictions were generated for all 58 consumer groups using the forward mixing model described in Section 2.1. Required inputs included basal isotope signatures (four primary producers and two detrital groups), diet proportions from the EwE model or the alternate diet configurations, and trophic discrimination factors (TDFs) for each isotope. A single TDF per stable isotope ratio was applied across all trophic transfers.

Monte Carlo simulation was used to propagate uncertainty in basal isotope signatures and TDFs. For each of 1000 simulation iterations, basal signatures were sampled from normal distributions informed by published means and variances (Table 2), with constraints imposed to preserve ecologically realistic ordering (e.g., phytoplankton δ¹³C depleted relative to macroalgae). Truncated normal distributions were used to maintain minimum separation between sources and prevent unrealistic overlaps in individual draws. Basal δ¹⁵N values were assumed to be indistinguishable within the model domain (Tarnecki and Patterson 2015). TDFs were independently sampled for each isotope (Δ¹³C ∼ N(1.5, 0.5), Δ¹⁵N ∼ N(3.0, 0.5), Δ³⁴S ∼ N(0.0, 0.5)) and applied uniformly across the food web (Post 2002; Post et al. 2007; Raoult et al. 2024). The selected distributions reflect commonly reported values while incorporating variability in trophic discrimination factors among taxa, diets, and ecological contexts (Caut et al. 2009; Wyatt et al. 2010; Stephens et al. 2023). Further details of sampling procedures can be found in supplementary materials (Supplementary Materials 1).

**Table 2.**
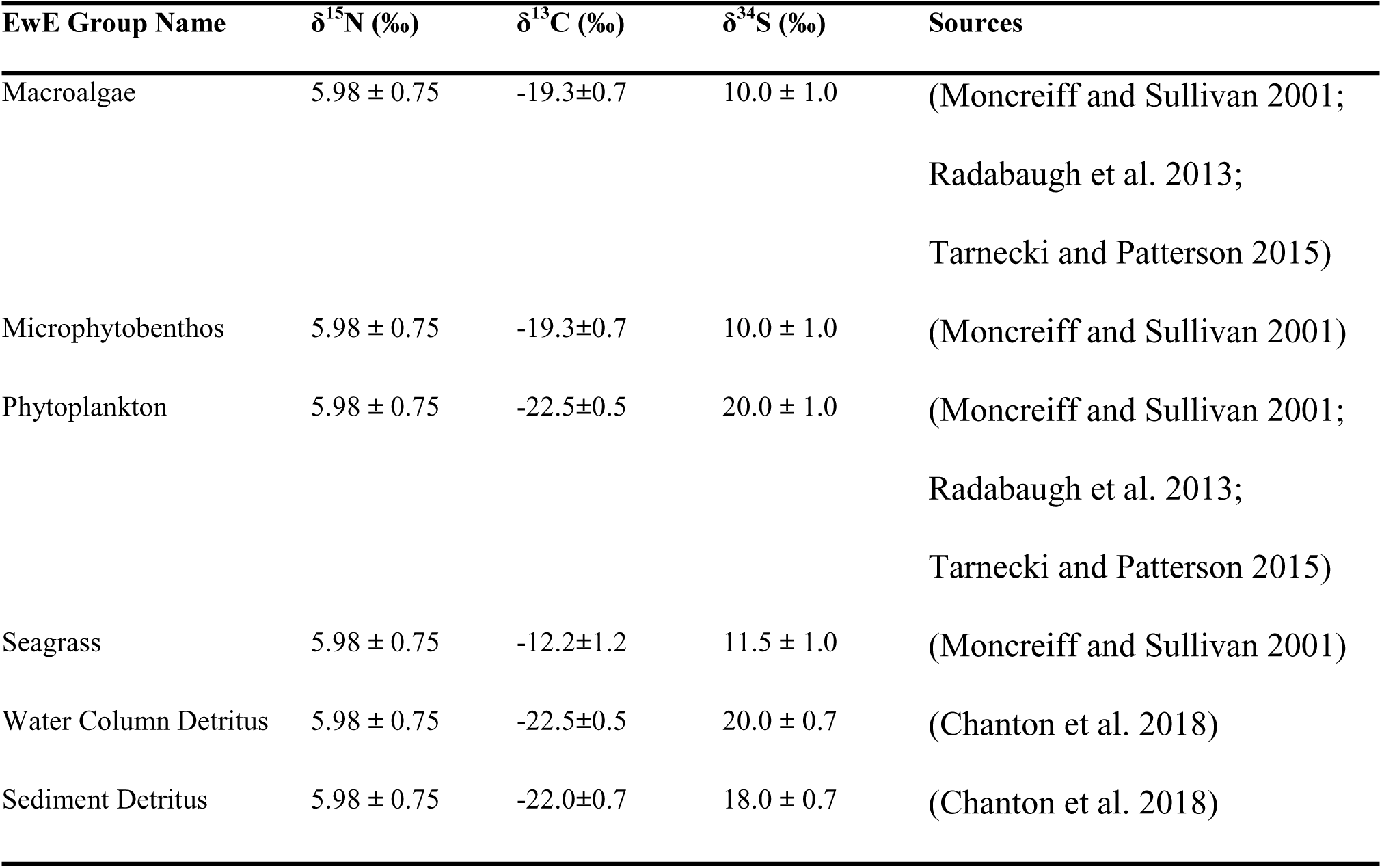
Basal source group δ^13^C, δ^15^N, and δ^34^S used for Monte Carlo simulation draws.

#### 2.2.4 Estimation of benthic–pelagic pathway contributions

Benthic and pelagic pathway contributions were estimated from predicted isotope values using δ¹³C and δ³⁴S (Equations 4-6). Representative benthic signatures were defined as biomass-weighted means of macroalgae and microphytobenthos, which dominate benthic production in the NGoM-EwE model, based on group biomass (B) parameters. Seagrass was excluded from benthic end-member calculations because its contribution to basal production was comparatively small in the NGoM-EwE model. Phytoplankton represented the pelagic basal member.

Equivalent representative values were calculated for δ¹³C. Trophic position and associated fractionation were accounted for using predicted δ¹⁵N values. Traditional inverse mixing model formulations were then applied to estimate pathway contributions for each consumer. Observed pathway contributions were calculated using the same approach applied to empirical isotope data (Equations 7-9). Residuals for pathway contributions were computed as the difference between observed and predicted benthic contributions (Δ = observed − predicted), where positive values indicate overestimation of pelagic pathway utilization.

#### 2.2.5 Model Scoring and Comparison

Model performance was evaluated using in-sample log predictive density (LPD), calculated by comparing observed isotope values with predicted distributions for each diet matrix configuration. Pointwise log predictive densities were summed across observations to quantify overall model fit, with higher values indicating better agreement. Observed isotope data were modeled using location–scale t-distributions to provide robustness to outliers and small sample sizes (Table 1). For groups with limited sample sizes (n = 2–3), the scale parameter was fixed using a pooled median absolute deviation (MAD) derived from groups with larger sample sizes. For groups with n ≥ 4, both location and scale parameters were estimated. A common degrees of freedom parameter (ν = 7) was used across all groups to ensure robustness to outliers.

Model performance was also evaluated using pathway-based metrics. For each consumer, the predictive distribution of benthic–pelagic contribution was compared with the observed value, and log predictive densities were summed across consumers to produce a pathway-level score.

These metrics were used to compare alternative diet matrices and assess their ability to reproduce both isotope signatures and inferred energy pathways.

#### 2.2.6 Drivers of Mismatch

To investigate potential drivers of mismatch, redundancy analysis (RDA) was used as a secondary diagnostic analysis to evaluate whether ecological traits, model structure or sampling characteristics explained variation between model predictions and stable isotope observations. The MLE diet matrix was selected for RDA because it provided a single statistically estimated diet configuration derived independently from the stable isotope data and served as the reference around which stochastic diet matrices were generated. Response matrices included LPD and standardized residuals for elemental (C, N, S) and pathway-level metrics (carbon- and sulfur-derived), each centered and scaled prior to analysis. Explanatory variables described ecological structure (trophic level, body size, omnivory, imported production, species richness), sampling characteristics (stomach and isotope sample size, data source composition, spatial extent, species coverage, coverage mismatch, and habitat maximum depth). Maximum depth was included as a proxy for habitat position along an inshore-offshore gradient. This variable was considered particularly relevant for δ³⁴S because benthic and pelagic sulfur signatures may become increasingly homogenized in shallow, turbulent environments, reducing pathway discrimination (Raoult et al. 2024).The EwE-derived omnivory index measures the variance in trophic levels among prey items (Christensen et al. 2004)and was used as a proxy for diversity of prey items in the diet.

Species coverage quantifies how many species of a NGoM-EwE functional group contained samples based on effective Shannon diversity (*S_eff_* = exp(*H*)) relative to total group richness. Coverage mismatch was calculated as the absolute difference in coverage between stomach contents (SCA) and stable isotope (SIA) datasets, |*Coverage_SCA,_* − *Coverage_SIA_*|, representing differences in how completely species within each functional group were sampled. This metric reflects the effective proportion of species represented within each group for a given data source. For single-species groups, coverage mismatch is zero when the species is represented in both datasets. For multi-species groups, mismatch increases when SCA and SIA data are derived from different subsets of species within the group, reflecting differences in within-group sampling completeness. RDA models were fit using the vegan package (Oksanen et al. 2026).

Multicollinearity was evaluated using variance inflation factors (VIF), and interpretation focused on the first two constrained axes. For visualization purposes, RDA figures display a subset of explanatory variables with among the largest biplot scores. Variables that were nearly collinear with stronger predictors were omitted to reduce redundancy and improve readability.

### 2.3 Software

Data analysis, simulation, and plotting were completed using R version 4.5.1 and the NGoM was accessed using Ecopath with Ecosim version 6.7.0.18858.

## 3. Results

### 3.1 NGoM-EwE base model predictions

Consumer isotope signatures were predicted for all consumers in the NGoM-EwE model (Figures 2–3). Comparison of simulated and observed consumer δ¹³C values indicated moderate mismatch; however, predictions from the forward mixing model were generally consistent with observed data. Most standardized residuals were within ±2 when accounting for uncertainty (Figure 5). Measured group means fell within Monte Carlo predictive intervals for all elements and for all 34 consumer groups with available data, with the exception of δ³⁴S for damselfish.

**Figure 2.**
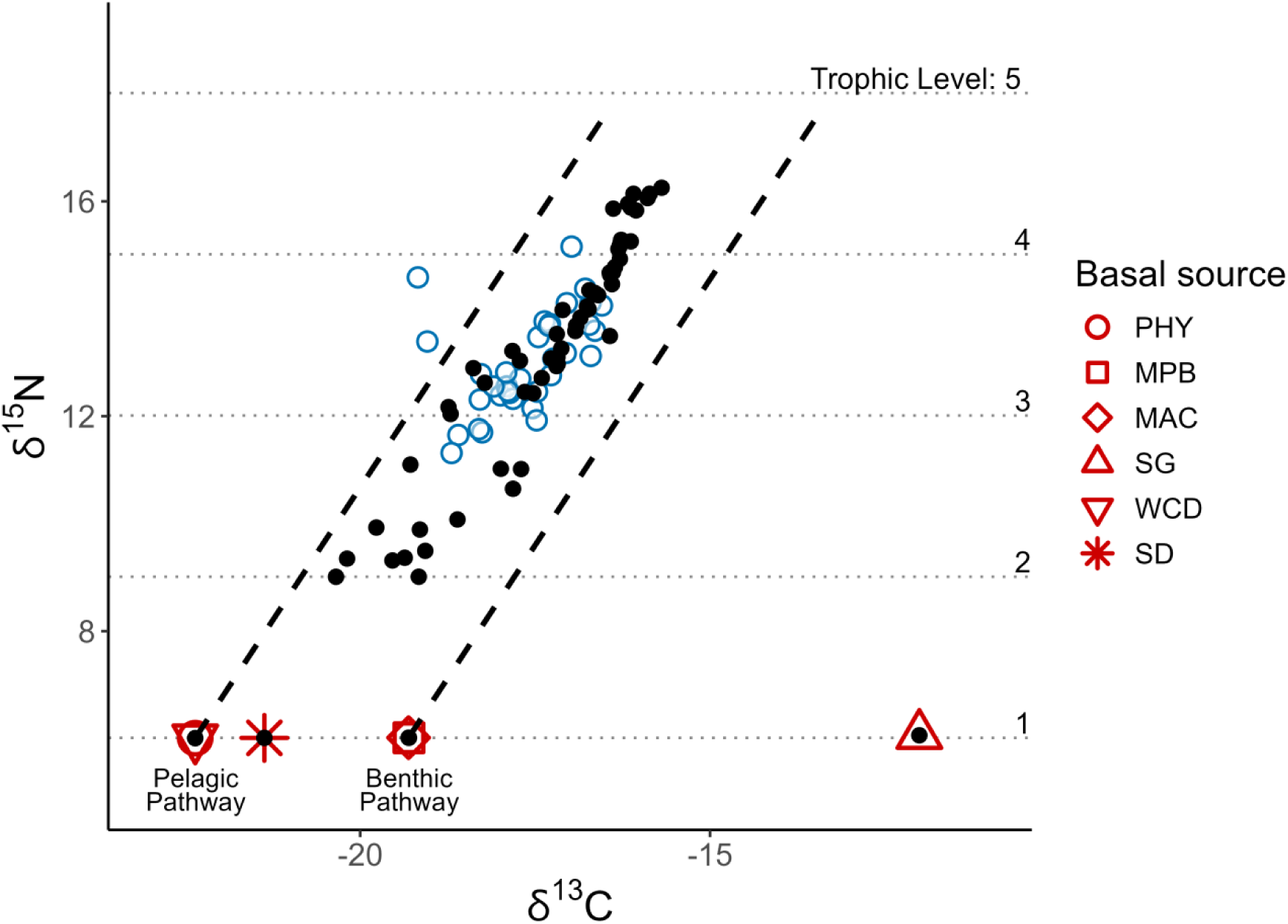
Observed (open blue circles) and predicted (filled black circles) fish muscle δ¹³C and δ¹⁵N signatures for the NGoM-EwE model. Predicted values are derived from forward mixing of diet matrices. Basal source endmembers (red symbols) and trophic enrichment boundaries are shown for reference. Dashed bands represent assumed mean trophic enrichment relationships, and horizontal lines denote trophic level increments. Agreement reflects consistency between modeled diet and isotope-implied trophic structure. Basal sources are phytoplankton (PHY), microphytobenthos (MPB), macroalgae (MAC), seagrass (SG), water-column detritus (WCD), and sediment detritus (SD).

**Figure 3.**
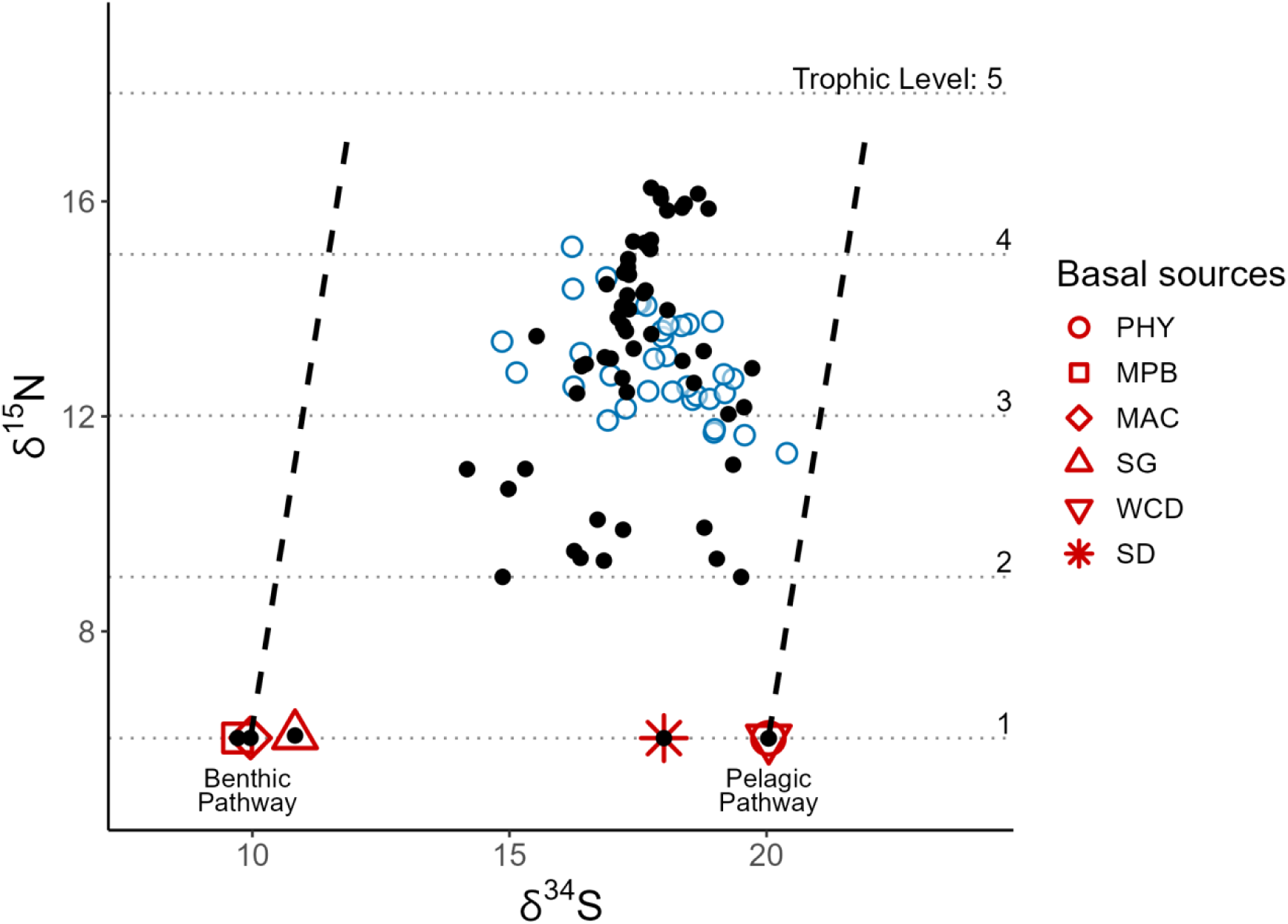
Observed (open blue circles) and predicted (filled black circles) consumer δ¹⁵N and δ³⁴S signatures for the NGoM-EwE model. Predicted values are derived from forward mixing of diet matrices. Basal source endmembers (red symbols) and trophic enrichment boundaries are shown for reference. Dashed lines represent assumed mean trophic enrichment relationships, and horizontal lines denote trophic level increments. Agreement between observed and predicted isotope signatures reflects consistency between modeled diets and isotope-implied trophic structure. Basal sources are phytoplankton (PHY), microphytobenthos (MPB), macroalgae (MAC), seagrass (SG), water-column detritus (WCD), and sediment detritus (SD).

**Figure 4.**
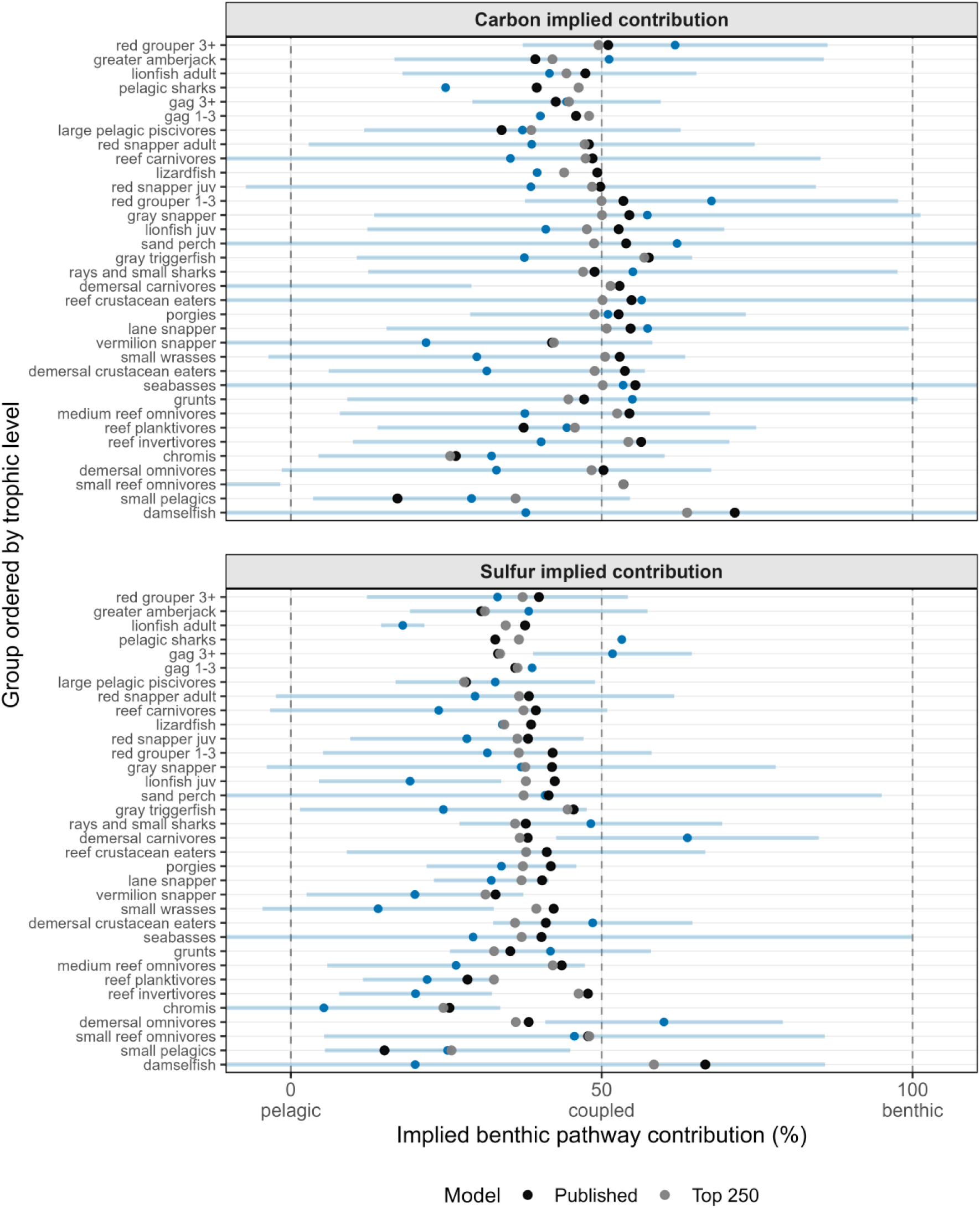
Benthic pathway contributions (%) for each functional group derived from the published model (black circles) and the mean of the top-performing stochastic ensemble (gray circles), compared with isotope-implied estimates and associated 95% confidence intervals (light-blue points and horizontal bars). Functional groups are ordered by trophic level. Panels display carbon- and sulfur-derived pathway estimates, with vertical dashed lines indicating pelagic-, coupled-, and benthic-dominated energy pathways. The stochastic ensemble represents the mean of the 250 highest-performing diet matrices generated through stochastic perturbation around the maximum-likelihood estimate.

**Figure 5.**
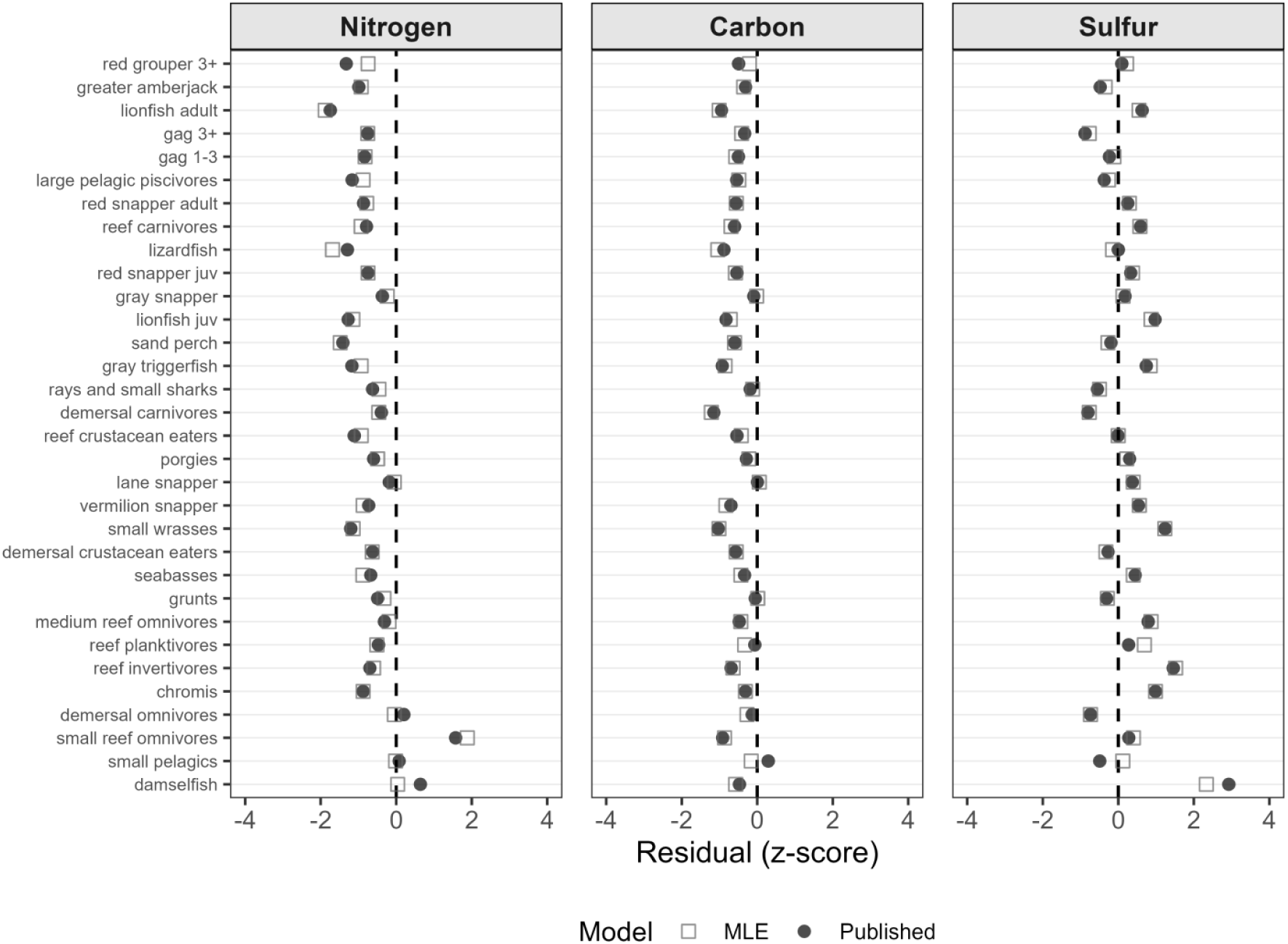
Standardized residuals (z-scores) for δ¹⁵N, δ¹³C, and δ³⁴S comparing predictions from the published NGoM-EwE model (filled black circles) and the maximum-likelihood diet matrix (MLE; open gray squares) across functional groups ordered by trophic level. The vertical dashed line indicates zero residual (perfect agreement), with points closer to zero indicating better agreement between model predictions and observed isotope values.

Standardized residuals for δ¹⁵N were negative for 30 of the 34 groups, indicating that modeled trophic levels were generally higher than those implied by empirical isotope data (Supplementary Table 1).

In contrast, inferred benthic pathway contributions revealed greater mismatch between modeled energy pathways and those implied by observed stable isotopes (Figure 6). For δ¹³C-based pathway estimates, 22 of 34 groups fell within the 95% predictive intervals generated by the Monte Carlo simulation. For δ³⁴S-based pathway estimates, only 4 of 34 groups fell within the corresponding predictive intervals (Supplementary Table 1), indicating substantial disagreement in sulfur-derived pathway representation. For 61.8% of groups, the direction of pathway misalignment was consistent between carbon- and sulfur-derived pathway estimates, with ∼62% of groups represented as relying too much on benthic energy when estimated via carbon and ∼71% when estimated via sulfur. Taken together, these results indicate that while the diet configurations in the NGoM-EwE model broadly reproduce observed stable isotope values, discrepancies remain in how energy pathways are represented through the diet matrix.

**Figure 6.**
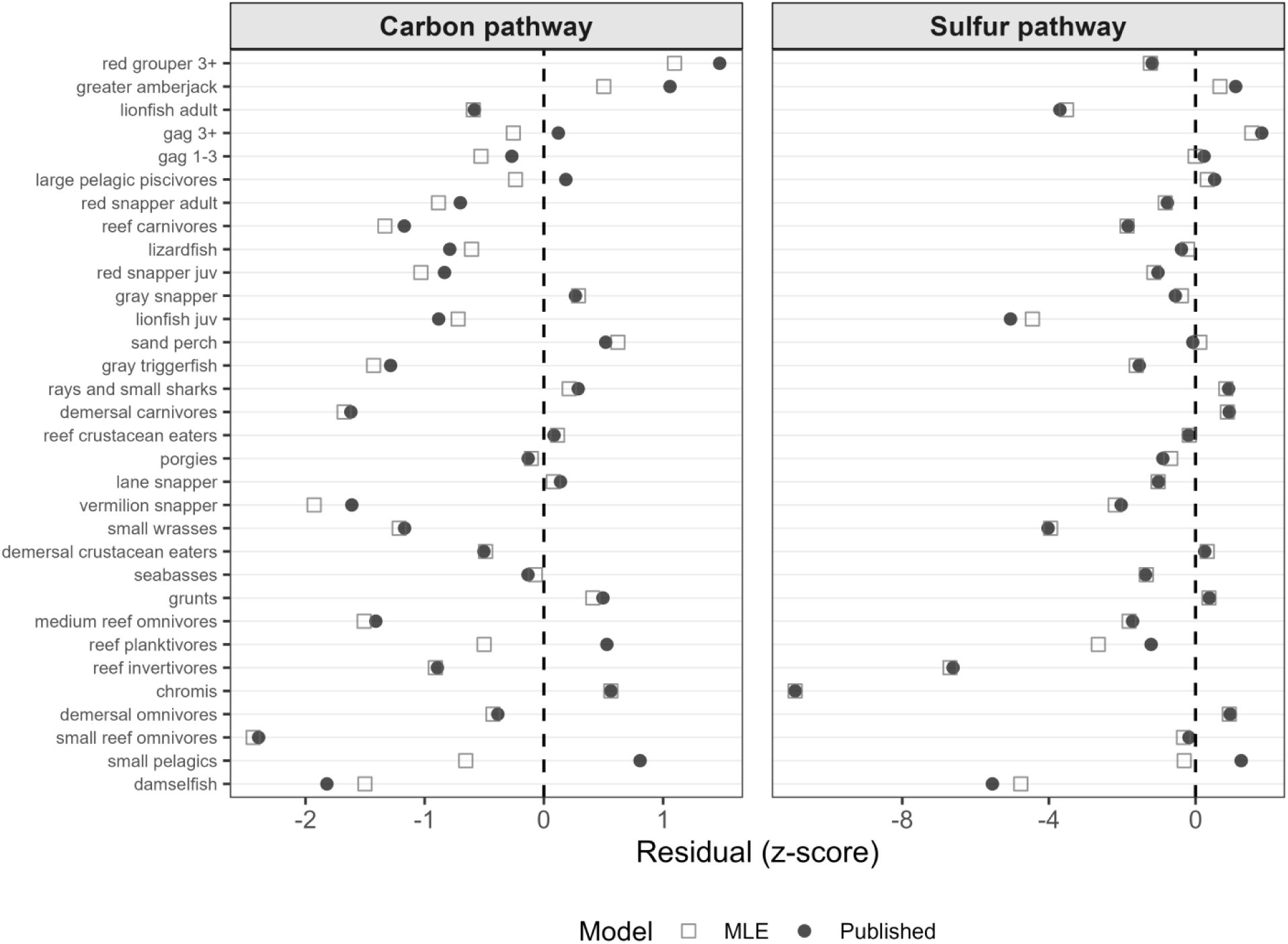
Standardized residuals (z-scores) of isotope-implied benthic pathway contributions derived from δ¹³C and δ³⁴S, comparing the published NGoM-EwE model (filled black circles) and the maximum-likelihood diet matrix (MLE; open gray squares) across functional groups ordered by trophic level. Panels show carbon- and sulfur-derived pathway estimates. The vertical dashed line indicates zero residual, corresponding to perfect agreement between modeled and isotope-implied pathway contributions. Residuals closer to zero indicate improved agreement.

### 3.2 Model comparison

All alternative diet matrix configurations outperformed the published, balanced diet matrix based on log predictive density (LPD) (Table 3). Improved model fit was reflected by less-negative LPD values and was consistent across total LPD, isotope-based LPD, and pathway-based LPD. The weighted average configuration produced better scoring, slightly higher combined LPD (−533.97) and isotope LPD (−177.89) compared to the maximum likelihood (MLE) configuration (−535.50 and −179.92, respectively). In contrast, the MLE configuration yielded a marginally higher pathway LPD (−355.57) than the weighted average configuration (−356.08). Model performance was modestly improved among the stochastically varied diet matrices, indicating that additional variation in diet proportions within empirically supported bounds, including for groups without direct isotope observations, can improve agreement with observed data. This finding suggests that, in addition to the choice of diet estimation method, incorporating uncertainty in diet proportions can further improve representation of trophic structure and energy flow.

**Table 3.**
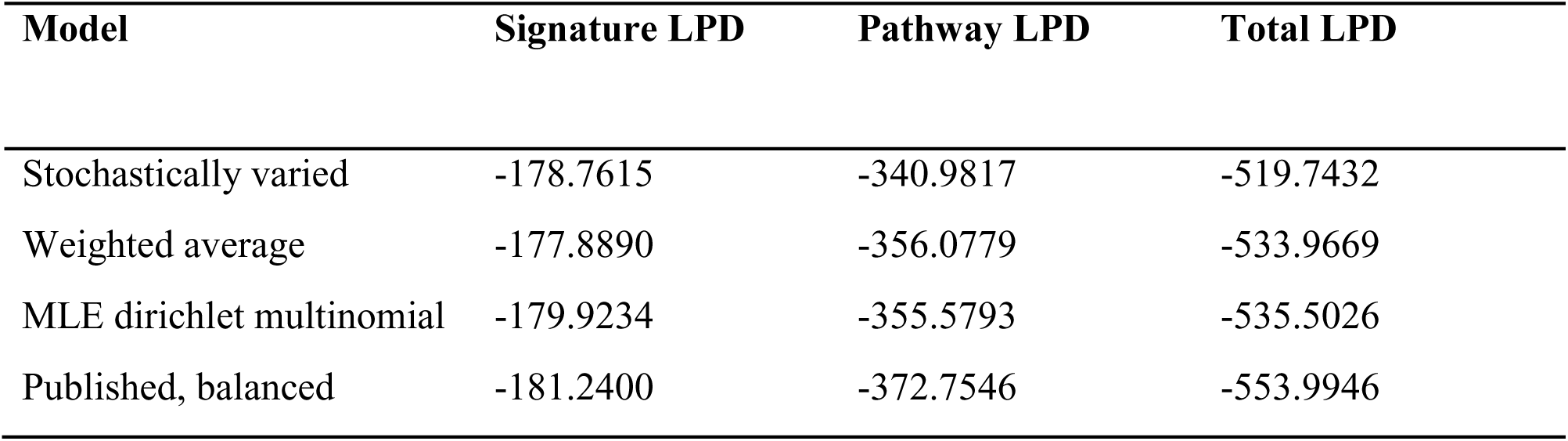
Comparison of log predictive densities (LPD) for published and alternative models. The Stochastically varied model reports the mean LPD across the 250 highest-scoring matrices (ranked by total LPD) from the 10,000 sampled configurations.

### 3.3 Mismatch in diet representation

Redundancy analysis (RDA) explained a substantial proportion of variation in both LPD values (48.9%; Figure 7) and standardized residuals (53.0%; Supplementary Figure 2). In the LPD model, variation was strongly structured along a primary axis (RDA1), which accounted for approximately 64% of the explained variance, while a secondary axis (RDA2) accounted for ∼19%. The primary axis was strongly associated with sampling and coverage variables, particularly the magnitude of coverage mismatch between stomach contents and stable isotope datasets, which emerged as the strongest individual predictor of diet representation mismatch. This axis also aligned with stomach sampling effort and group-level species richness, indicating that differences in species coverage within functional groups define the dominant gradient in model performance.

**Figure 7.**
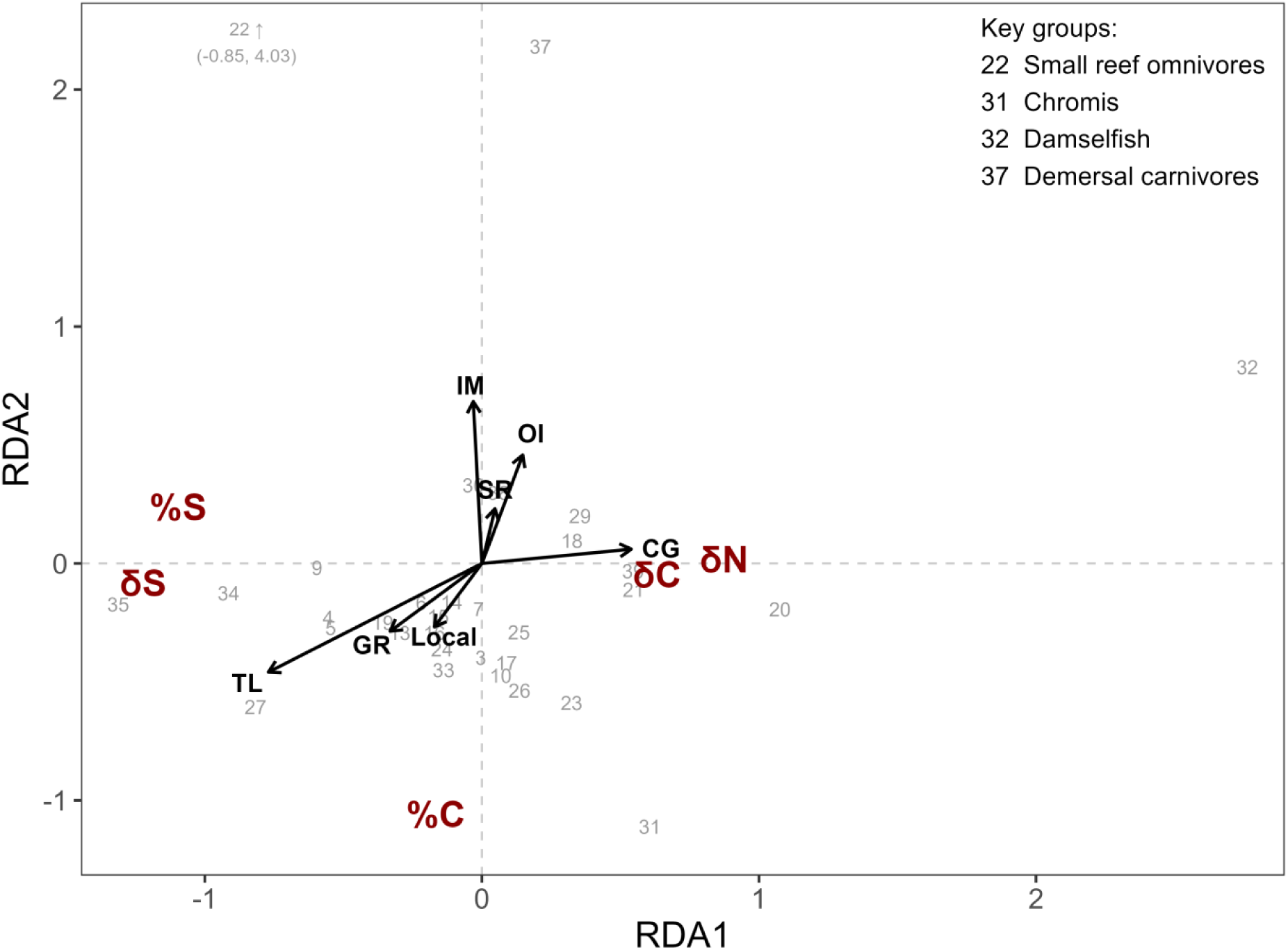
Redundancy analysis (RDA) of log predictive density (LPD) metrics for δ¹³C, δ¹⁵N, δ³⁴S, and implied benthic pathway contributions (%C and %S), showing relationships between model-data agreement and ecological and sampling variables. Points represent functional groups, and vectors indicate explanatory variables, with direction and length reflecting their association with model fit. Explanatory variables shown are trophic level (TL), omnivory index (OI), species richness (SR), imported diet fraction (IM), sampling coverage gap (CG), geographic range of SIA samples (GR), and proportion of diet samples collected specifically for this study rather than obtained from repositories (Local). Functional groups are identified by numerical codes corresponding to group names listed in Table 1; selected influential groups are identified directly in the figure.

The second axis was structured by ecological variables, including trophic level, omnivory, and imported production. Higher values of the EwE-derived omnivory index and imported production fractions opposed higher trophic level and size-related variables, suggesting a gradient in ecosystem openness and trophic organization. In contrast, variation in residuals was more evenly distributed across ordination axes, with weaker dominance of the primary axis.

Although explanatory variables accounted for a similar proportion of total variance, no single gradient strongly structured residual patterns, indicating the absence of a consistent directional bias in model error.

## 4. Discussion

### 4.1 Forward isotope simulation as a validation tool

The forward isotope simulation framework provides a new approach for evaluating trophic interactions in ecosystem models by directly linking diet matrices to predicted stable isotope signatures. A key advantage of the framework is that it evaluates trophic structure directly in isotope space rather than through derived summary metrics. Traditional comparisons of model output to stable isotope data often rely on reduced metrics such as trophic level (δ¹⁵N) or omnivory indices, which collapse complex trophic interactions into single values (Navarro et al. 2011; Deehr et al. 2014; Pedersen 2022). This simplification is especially necessary in multi-source systems, where isotope mixing models cannot uniquely resolve contributions from multiple prey or basal sources and are often reduced to summary metrics (Stock et al. 2018).

While these metrics are useful, they inherently obscure the underlying mixing structure of energy pathways and basal source contributions. By directly predicting stable isotope signatures, the forward approach preserves the multivariate structure of isotope space and the constraints imposed by multiple tracers and food web structure. As a result, the framework can be applied consistently across simple and complex systems without reducing trophic interactions to simplified metrics.

Study results illustrate the advantage of a forward modeling approach. Traditional inverse isotope mixing models seek to estimate diet composition from consumer isotope signatures and can become indeterminate when multiple prey combinations produce similar isotopic solutions (Smith et al. 2013; Stock et al. 2018). In contrast, the framework presented here uses the diet matrix as a starting point and evaluates whether the resulting isotope predictions are consistent with observed data. Consequently, model evaluation is not contingent on uniquely resolving source contributions. Because all tracers are propagated through the same network mixing framework, δ¹³C, δ¹⁵N, and δ³⁴S can be evaluated consistently without reducing each isotope to separate ecological indices or validation metrics. This reduces the need for tracer-specific assumptions and allows direct assessment of whether a proposed trophic structure is compatible with the available isotope data. However, the framework does not identify a unique "correct" diet configuration. Rather, model scores quantify the plausibility and internal consistency of a proposed diet matrix given the isotope observations.

### 4.2 Insights from the NGoM case study

Predicted fish muscle stable isotope values showed generally good agreement with observations, consistent with previous comparisons of δ¹⁵N-derived trophic position and EwE estimates (Kline Jr and Pauly 1998; Deehr et al. 2014; Pedersen 2022). In particular, agreement in δ¹⁵N suggests overall trophic positioning is reasonably captured. However, the strong mismatch observed in benthic–pelagic pathway estimates, particularly for δ³⁴S, demonstrates that models can reproduce observed isotope signatures while still misrepresenting how energy flows through the food web. Stable isotope analyses have previously revealed shifts in trophic ecology and resource use following environmental disturbance that were not readily apparent from stomach contents data alone (Tarnecki and Patterson 2015), as well as broader alterations in ecosystem energy pathways associated with invasion and restoration (Dibble and Meyerson 2014). The pathway-level discrepancies identified here similarly suggest that stable isotopes can provide information on food-web structure and energy flow beyond that contained within stomach contents-derived diet matrices.

The NGoM case study is particularly well suited for evaluating benthic–pelagic coupling due to clear separation in basal isotope signatures. This separation allows pathway contributions to be inferred independently from δ¹³C and δ³⁴S, providing multiple lines of evidence for evaluating energy flow (Dahl and Patterson 2014; Tarnecki and Patterson 2015). The consistent tendency for the model to overrepresent benthic energy contributions suggests that pathway imbalance may be embedded within the diet parameterization. Because benthic and pelagic production pathways support different components of marine food webs and can influence ecosystem structure and responses to disturbance (Dibble and Meyerson 2014; von Biela et al. 2016), misrepresentation of their relative contributions may affect ecological interpretation and model projections. Consequently, misrepresentation of these pathways may affect predictions of ecosystem responses to disturbance, habitat change, and other environmental stressors. Correctly representing energy pathways is therefore important not only for model realism, but also for generating reliable ecological forecasts and management advice.

Although the consistent direction of pathway misalignment across carbon- and sulfur-based estimates suggests that at least part of the observed discrepancy reflects systematic bias in diet representation rather than random error alone, several ecological mechanisms could also contribute to these discrepancies and may reflect ecological complexity in benthic-pelagic coupling rather than deficiencies in the Ecopath diet matrix alone (Kopp et al. 2015; Wang et al. 2020). Consumers associated with benthic habitats do not necessarily rely exclusively on benthic energy pathways (von Biela et al. 2016). For example, site-attached reef fishes may occupy benthic or demersal habitats while feeding extensively on pelagic prey, resulting in isotope signatures that integrate energy from multiple pathways. Similarly, sulfur signatures of benthic primary producers may not always reflect sediment-derived sulfur sources. Macroalgae can assimilate nutrients directly from the water column, potentially reducing the expected separation between benthic and pelagic δ³⁴S signatures (Moncreiff and Sullivan 2001). In addition, benthic detrital pools are often supported by sinking particulate organic matter (marine snow), causing sediment detritus to retain relatively pelagic isotopic characteristics and further narrowing the distinction between benthic and pelagic sulfur pathways (Chanton et al. 2018). Together, these processes can blur pathway distinctions inferred from isotopes and contribute to disagreement between modeled and isotope-implied pathway contributions even when overall isotope values are reproduced accurately. While δ³⁴S provided valuable additional resolution in this study, its usefulness for model evaluation will vary among ecosystems and should be assessed on a case-by-case basis (Raoult et al. 2024).

### 4.3 Drivers of model-data mismatch

The NGoM-EwE model represents a relatively data-rich system, making the strong influence of species coverage mismatch particularly notable. The redundancy analysis identified species coverage mismatch between stomach contents (SCA) and stable isotope (SIA) datasets as the strongest observed association with model-data mismatch. This indicates that differences in which species are sampled, and how completely they are represented within functional groups, can have a greater influence on model performance than sampling effort alone.

In EwE models, functional groups often aggregate multiple species, while isotope data typically represents only a subset of those species. In this study, most non-zero coverage mismatches occurred where SIA data represented fewer species than SCA data, suggesting that isotope observations may not always represent the same species composition represented by the functional group and underlying diet data. As a result, apparent disagreement between predicted and observed stable isotope signatures may arise even when the underlying diet matrix accurately represents trophic relationships within the functional group. This demonstrates that mismatch between model predictions and isotope observations may reflect sampling artifacts rather than structural flaws in the model itself. For isotope-based validation of aggregated functional groups, representativeness may be as important as sample size. Isotope samples must capture the species composition implied by the functional groups, rather than overrepresent populous or easily sampled species. Without alignment in species coverage between datasets, comparisons between modeled diets and isotope observations may be biased, particularly when evaluating pathway-level metrics. As a result, coordinated collection of stomach contents and stable isotope samples from the same species assemblages may be more valuable for model validation than simply increasing sample sizes within a single data source.

In addition to differences in species coverage, mismatch increased with trophic level, indicating that uncertainty associated with diet composition, trophic discrimination factors, and basal isotope signatures propagates through the food web. Errors in lower trophic levels accumulate through predation, leading to greater discrepancies at higher trophic positions (Ainsworth et al. 2010; Trijoulet et al. 2019). Although uncertainty in trophic fractionation and baseline values was incorporated through Monte Carlo simulation, the observed increase in mismatch with trophic level suggests that uncertainty in diet composition may still be underestimated. While imported prey fractions were relatively small in this case study and were excluded from analysis, their presence highlights a limitation of the framework. Additional uncertainty may also arise from incomplete representation of consumer diets within the model boundary, as higher trophic level groups in the NGoM-EwE model included greater proportions of imported diet. For highly migratory species or systems with substantial external inputs, interpretation of isotope predictions may therefore require additional caution.

### 4.4 Implications for diet matrix parameterization

Comparison of alternative diet matrices demonstrated that both diet estimation methods and uncertainty in prey proportions influence the agreement between modeled diet and diet implied by stable isotope data. Diet matrices derived directly from SCA data showed closer agreement with isotope-implied diet than the published, balanced model, indicating that adjustments introduced during model balancing may degrade representation of trophic interactions. Care should therefore be taken when modifying diet matrices during model balancing, and independent data sources such as stable isotopes, coupled with validation frameworks like the one described here, may provide a useful check on the ecological realism of balancing decisions.

Despite limited data for lower trophic level groups, stochastic variation in invertebrate diets improved model performance. Invertebrate diets are often poorly characterized in ecosystem models and are frequently borrowed from other regions, species, or previously published models. The sensitivity observed here suggests that improving diet information for lower trophic level groups may yield disproportionate benefits for ecosystem model performance. These patterns reinforce the importance of evaluating trophic structure at the network level, where errors at lower trophic levels can influence predictions across the entire food web.

Further improvements achieved through stochastic variation in prey proportions support the importance of explicitly representing uncertainty in diet composition (Masi et al. 2014; Steenbeek et al. 2018). Fixed diet matrices may underestimate variability in trophic interactions, whereas incorporating uncertainty allows for more realistic representation of energy flow. The improved agreement achieved by stochastic diet matrices demonstrates that multiple plausible diet configurations can produce meaningfully different isotope predictions and interpretations of energy flow, even when constrained by the same empirical data. Ecosystem models should therefore explicitly evaluate the sensitivity of model outcomes to uncertainty in diet composition and consider alternative parameterizations when sufficient dietary information is available.

Because the framework evaluates agreement rather than identifying a unique solution, care must be taken to avoid overfitting diet matrices to isotope data. Stable isotope observations should be treated as complementary constraints alongside stomach contents data, rather than as a substitute for them. In this study, stochastic variation was constrained using empirically derived bounds from SCA data, ensuring that alternative diet configurations remained biologically plausible. This approach provides a practical framework for integrating multiple dietary data sources while maintaining ecological realism.

### 4.5 Modeling assumptions and sources of uncertainty

Several modeling assumptions should be considered when applying this framework. Basal source signatures remain a key source of uncertainty. Although incorporated through Monte Carlo simulation, shifts in baseline values can influence both predicted isotope distributions and interpretation thresholds. Geographic range of isotopic sampling contributed to variation in model performance, suggesting that spatial variability in isotopic baselines may influence comparisons between modeled and observed isotope values. This consideration may be particularly important for large model domains that span environmental gradients.

The use of a single trophic discrimination factor (TDF) across groups represents a simplifying assumption. TDFs are known to vary among taxa, diets, trophic positions, tissues, and ecological contexts (Caut et al. 2009; Wyatt et al. 2010; Stephens et al. 2023). Incorporating group-specific or diet-specific TDFs may improve predictions, particularly for microbial pathways or systems with heterogeneous trophic processes. Although not implemented in this case study, the matrix formulation allows for group-specific TDFs to be applied at the level of individual rows in the system. Where improved estimates of TDFs are available, they can be readily incorporated within this framework.

### 4.6 Future directions

This framework extends stable isotope analysis from individual consumers to network-wide inference. Importantly, ecosystem models can generate isotope predictions for all functional groups within a food web, including groups for which stable isotope observations are unavailable. This capability is particularly valuable because isotope datasets are often incomplete, even in relatively data-rich systems such as the NGoM case study and may provide a means to extend inference beyond directly sampled taxa. Although implemented here as a static simulation, the framework provides the baseline isotope configuration from which dynamic simulations of isotope signatures can be initialized.

Dynamic isotope modeling offers the potential to capture temporal variability, isotopic turnover, and ecosystem responses to environmental change. While dynamic mixing models are being developed independently of ecosystem models (Ballutaud et al. 2022), coupling ecosystem model output with stable isotope flux models has also been demonstrated (Rastetter et al. 2005), but remains underutilized. Coupling forward isotope simulation with dynamic ecosystem models would enable prediction of time-varying isotope signatures throughout the food web, allowing direct comparison with observational datasets and providing a powerful tool for ecosystem monitoring, model validation, and ecological forecasting.

## 5. Conclusions

This study presents a reproducible framework for evaluating trophic structure in ecosystem models using stable isotope data. Results from the NGoM-EwE case study demonstrate that models may reproduce isotope signatures while misrepresenting underlying energy pathways, emphasizing the importance of pathway-based diagnostics. Within-group data coverage mismatch, particularly differences in species coverage between stomach contents and isotope datasets, emerged as a primary driver of model error, highlighting a critical consideration for sampling design.

Importantly, this framework does not identify a single “correct” trophic structure but instead evaluates the consistency of a given diet matrix with observed isotope data. Model scores should therefore be interpreted as measures of plausibility and internal consistency between data sources rather than definitive estimates of diet. By combining forward isotope modeling, diagnostic metrics, and model scoring, the framework provides a practical tool for evaluating and improving the representation of trophic interactions in ecosystem models.

## Author Contributions

**Megan Siemann**: Methodology, Formal Analysis, Writing - Original Draft. **Kieran Tierney:** Writing - Review & Editing and Validation. **Will Patterson**: Investigation, Resources, Project Administration. **David Chagaris**: Supervision, Writing - Review & Editing.

## Data Availability

Full code is available (github. Previously published data used are publicly available through GRIIDC repository (https://data.gulfresearchinitiative.org; https://doi.org/10.7266/N7445JW6; https://doi.org/10.7266/N7JM27KV). The full datasets are available upon request.

## Declaration of generative AI and AI-assisted technologies in the manuscript preparation process

During the preparation of this work the authors used Microsoft copilot to correct grammar and improve clarity of language. The authors reviewed and edited the resulting content and take full responsibility for the content of the published article.

## Acknowledgements

Thanks are given to Matt Campbell and Laura Reynolds both of whom provided guidance and feedback during preliminary stages of the project.

