## Supplementary Materials for "Validating static food web model diet matrices using stable isotope simulation"

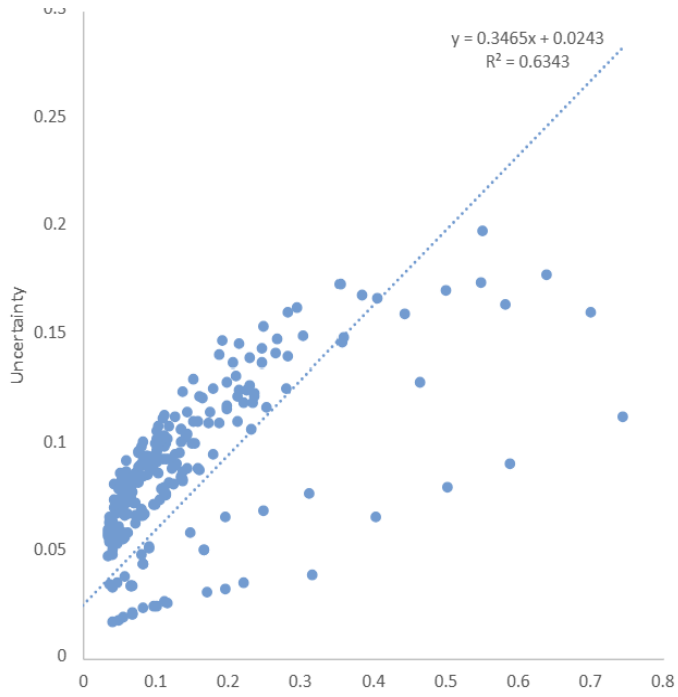


**Supplementary Figure 1.** Relationship between prey proportion and uncertainty for NGoM-EwE consumer diets estimated using a Dirichlet–multinomial model. The fitted linear relationship was used to assign pooled uncertainty bounds for invertebrate diets prior to stochastic variation of prey proportions.

**Supplementary Material 1.** Monte Carlo draws of basal source signatures:

Carbon ($\delta^{13}C$)

Carbon basal signatures were generated to preserve strong separation between benthic and pelagic sources while allowing correlated variability among benthic producers. Macroalgae was treated as the primary reference distribution (normal, mean −19.3‰, SD 0.7), and microphytobenthos (MPB) was modeled as a tightly scaled version of macroalgae (±~2.5%), reflecting similar isotopic composition. Phytoplankton was constrained to be isotopically lighter than benthic sources, with a minimum separation of 0.5‰ enforced via truncated normal draws incorporating combined variance. Seagrass was modeled as isotopically heavier than macroalgae, with a minimum separation of 2‰ and larger variance to reflect greater reported variability. Water column detritus was assumed to closely track phytoplankton with minimal additional noise, while benthic detritus was constrained to be equal to or slightly enriched relative to pelagic detritus. A trophic discrimination factor (TDF) for carbon was drawn from a normal distribution (mean 1.5‰, SD 0.5).

Nitrogen ($\delta^{15}N$)

Nitrogen signatures were modeled with a shared baseline to reflect relatively limited separation among basal sources in $\delta^{15}N$ space. A common baseline distribution (mean 5.98‰, SD 0.75) was first drawn, and individual sources (macroalgae, MPB, phytoplankton, and detrital pools) were generated as small deviations around this shared value (SD ~0.25). Seagrass was treated as slightly enriched relative to other sources, with a small offset and increased variance to reflect broader reported variability. This approach preserves strong covariance among sources while allowing realistic variability. Nitrogen trophic discrimination factors were drawn independently from a normal distribution (mean 3‰, SD 0.5), representing typical enrichment per trophic transfer.

Sulfur ($\delta^{34}S$)

Sulfur basal signatures were constructed to maximize contrast between benthic and pelagic pathways while maintaining realistic relationships among sources. Macroalgae was modeled as the primary benthic reference (normal, mean 10‰, SD 1), and MPB was generated as a slightly depleted but tightly constrained proportion of macroalgae. Phytoplankton was required to be isotopically enriched relative to benthic sources, with a minimum separation of 0.5‰ enforced using truncated normal draws with combined variance. Seagrass was allowed to vary around benthic values with broader bounds (−1.5‰ to +3‰ relative to macroalgae), reflecting more variable sulfur sourcing. Water column detritus closely tracked phytoplankton, while benthic detritus was constrained to be equal to or slightly depleted relative to pelagic detritus. Sulfur trophic discrimination factors were drawn from a normal distribution (mean 0.5‰, SD 0.5), reflecting minimal fractionation.

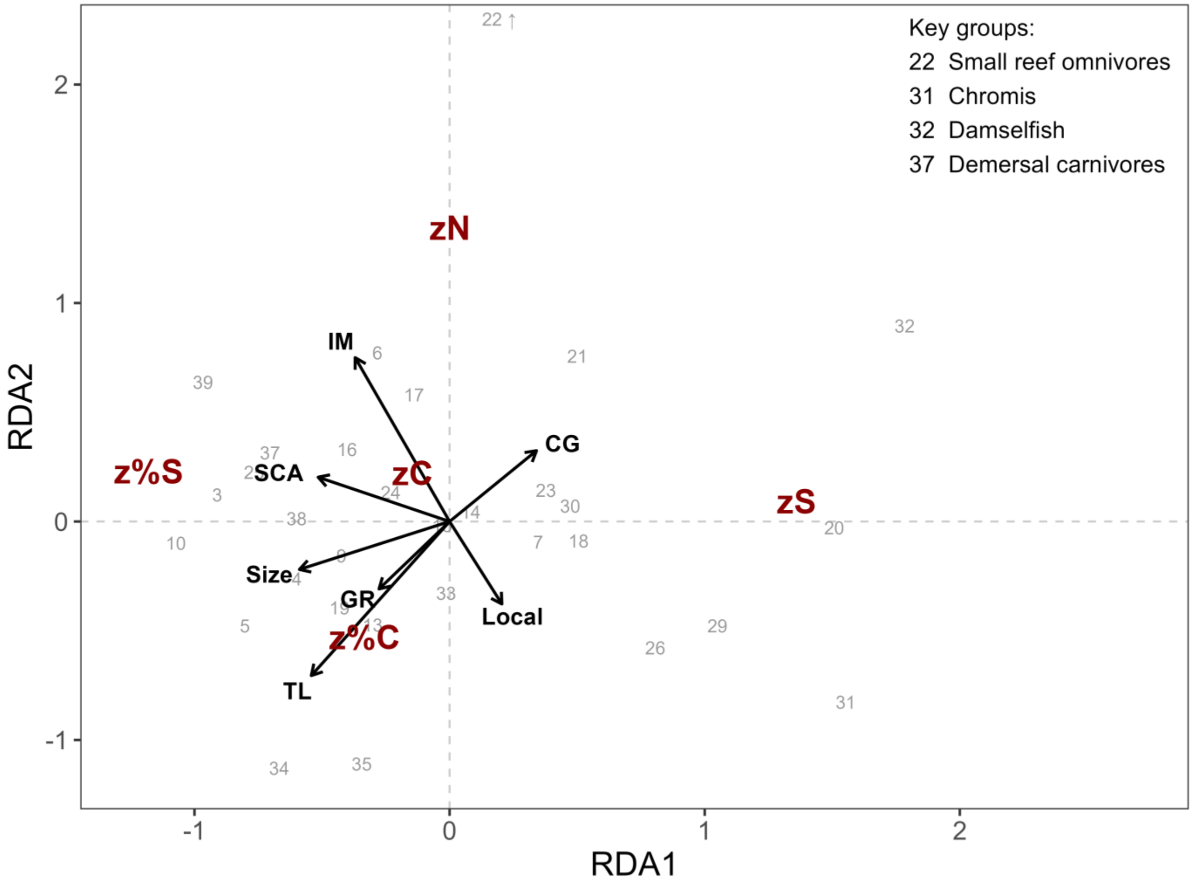


**Supplementary Figure 2.** Redundancy analysis (RDA) of standardized residuals (z-scores) for $\delta^{13}C$, $\delta^{15}N$, $\delta^{34}S$, and implied benthic pathway contributions (%C and %S), showing relationships between model-data mismatch and ecological and sampling variables. Points represent functional groups, and vectors indicate explanatory variables, with direction and length reflecting their association with residual structure. Explanatory variables shown are trophic level (TL), imported diet fraction (IM), observed body size (Size), diet sampling effort (SCA), sampling coverage gap (CG), proportion of diet samples collected specifically for this study rather than obtained from repositories (Local), and geographic range of SIA samples (GR). Functional groups are identified by numerical codes corresponding to group names listed in Table 1; selected influential groups are identified directly in the figure.

**Supplementary Table 1.** Table 3. Predicted stable isotope values by EwE group (mean, σ) and predicted stable isotope values, mean (95% predictive interval) from Monte Carlo Simulation. See separate excel document.
